# Invariant set theory for predicting failure of antibiotic cycling

**DOI:** 10.1101/2024.02.06.579227

**Authors:** Alejandro Anderson, Matthew W. Kinahan, Alejandro H. Gonzalez, Klas Udekwu, Esteban A. Hernandez-Vargas

## Abstract

The limited availability of antibiotics and the need for prompt decision-making present significant challenges for healthcare practitioners. When faced with this situation, practitioners must prioritize their approach based on several key factors. By leveraging the emergent understanding of collateral sensitivity among antibiotic-exposed pathogens, we demonstrate the utility of control invariant sets to predict treatment failure when antibiotic cycling is applied as a therapeutic strategy aiming to eradicate or prevent emergence of multi-drug resistant pathogens. Our results here pave the way for point-of-care diagnostic technologies to identify infections and select appropriate treatments quickly, reducing unnecessary antibiotic use.

## 1 Introduction

Antimicrobial drug resistance (AMR) was uncovered in the 1950s and it is well known that the deployment of antimicrobial without prescription has increased the AMR crisis [1, 2, 3, 4, 5, 6]. AMR has mainly been addressed by the discovery of new classes of antibiotics but it was reported that the development of antimicrobials might not keep up with the escalation of AMR in the next decades [7, 8, 9]. In addition, the combination of drugs necessary to treat AMR has triggered the emergence of multidrug-resistant (MDR) infections [10, 11, 12, 13] further aggravating the AMR crisis. Accordingly, control strategies that extend the life span of existing antibiotics and reduce the MDR are currently being studied [14, 15, 16, 17].

One approach has been focused on exploiting synergistic and antagonistic effects between antimicrobials [18], like orthogonal sensitivities that arise from the evolution to resistance which is known as collateral sensitivity (CS). CS was first described for Escherichia coli in 1952 [19] and it was observed in microbes infections and in cancer as well [20]. The CS phenomenon occurs when the genetic changes accrued on the develop of resistance towards one agent simultaneously decrease fitness to a second agent [21] and it is thought to be caused by a trade-off of pleiotropic resistance mutations [22]. More importantly, CS can be used to design drug combinations therapy: for instance by including mutual CS drugs in drug-cycling regimen [23, 24, 25]. To further understand the extent of CS, laboratory evolution has been deployed to map out CS profiles between the known antibiotics for specific drug-resistant strains [26, 27, 28, 29]; genome sequencing to characterize the components that contribute to CS [30, 31, 32, 33]; and the development of mathematical modeling that provides insight into the collateral dynamics network and supports control strategies to reduce the growth of MDR [34, 35].

Mathematical modeling establishes a high level of control involving factors such as pharmacodynamic (PD) and pharmacokinetic (PK) [36], optimal time of drug exposure [37], between others with relevance on the effects antibiotics have on bacterial growth, inhibition, killing, and mutation [38, 39, 40, 41]. Regarding collateral effects, mathematical models have been developed to evaluate *sequential drug regimens* in vitro and silico [42, 43], to assess the *robustness* of collateral sensitivity [43], to exploit its statistical structure and designing optimal policies [44], to evaluate possible *reversion of evolution towards resistance* [45] and to compare *cycling vs mixing* treatments [46]. None of these models however tackled the CS phenomenon with a combinatorial mutation network which has already been used for multidrug resistance (without collateral effects) on cancer [47], for which it was assumed that a mutation that confers resistance to one drug does not confer resistance to any of the other drugs in use; framework that leads to a dead-end towards MDR. The epistatic interactions between genes involved in CS is a complex and not fully understood phenomenon but some magnitudes and directions affected by this phenomenon can be measured and framed within combinatorial mutation networks and - by dynamic network-based analysis [48] and the set-control theory [49] - we can determine drug combinations that contain the infection.

Eliminating an infection requires driving the system states to the healthy equilibrium at the origin. This is referred to as the regulation problem in control [50, 51, 52]. However, stabilizing the origin is not always feasible, especially in the context of evolutionary dynamics that lead to resistance. In such cases, more flexible control goals, such as invariant sets, become invaluable [53, 54]. Invariant sets, unlike fixed equilibria, enhance controllability by offering alternative stable regions and they provide safety zones to maintain infection suppression [55]. Such invariant sets play a fundamental stabilization role as the only regions formally stabilizable under Lyapunov analysis [56]. In this study, we introduce a combinatorial mutation network that models qualitative collateral sensitivity data for the assessment of sequential drug regimens. We employed a switched system framework [57, 58], extensively utilized in biomedical control problems [59, 60, 58] and recently adapted for sequential drug regimens with mutation dynamics [**?**, 61], with the goal of targeting phenotype states associated with specific collateral effects to improve treatment success in chronic infections. The model suggests that collateral sensitivity profiling can forecast the emergence and proliferation of MDR strains. Furthermore, we explore invariant regions for resistance evolution population trajectories that determine the conditions under which mutual collateral sensitivity possesses the capacity to contain a chronic infection.

## 2 Mathematical abstraction

The European Committee on Antimicrobial Susceptibility Testing (EUCAST) defines a microorganism susceptibility to a level of antimicrobial activity as a high likelihood of therapeutic success. Conversely, resistance is defined as a high likelihood of therapeutic failure. These categories are formally characterized by breakpoints determined in standard phenotype test systems [62, 63]. Ideally, clinical breakpoints distinguish between patients who are likely or unlikely to respond to antimicrobial treatment.

For convenience, the minimum inhibitory concentration (MIC) can be used as a parameter of antibiotic action [38], since it was reported a linear relationship between an increase in the MIC and hospital moralities [64]. It is defined as follows:

### Definition 1

(Minimum Inhibitory Concentration - MIC). *The MIC breakpoint is defined as the lowest concentration of a drug that inhibits the visible growth of a microorganism*.

*Collateral sensitivity* between drugs *A* and *B* can be defined as a decrease in the MIC of *B* for the antibiotic-resistant strain, when expose to drug *A*. Conversely, an increase in the MIC defines a *collateral (or cross) resistance*. On the other hand microbiology laboratories uses clinical breakpoints to categorize microorganisms as clinically susceptible or resistant dependent on the quantitative antimicrobial susceptibility as indicated by the MIC value [65, 66]. Figure 1 illustrates the scenario for two hypothetical drugs *A* and *B*, for the case of combining antibiotics in sequential order (not for mixtures drugs). According to Figure 1 the MIC of the wild-type (Variant *V* 0) indicates a profile of susceptibility to both drugs, after *V* 0 is expose to drug *A* the MIC increase respect to *A* and respect to *B*, i.e. drug *A* shows cross resistance. On the other hand, after *V* 0 is expose to drug *B* the MIC increase respect to *B* but it decrease respect to *A*, i.e. drug *B* shows collateral sensitivity. To determine the susceptible/resistant profile of emerging variants following drug exposure, collateral sensitivity effects can is formalized in next section using a combinatorial mutation network and, then linked the network with a switched system to model the population trajectories under drug sequential therapies.

**Figure 1.**
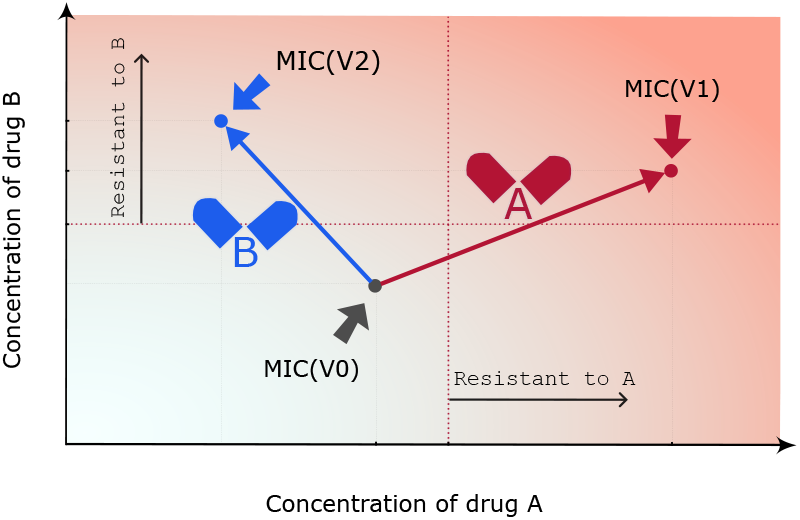
Drug-driven vectors describe the direction the MIC evolves under drug exposure. The drug-driven vector of *A* (red) shows cross-resistance and the vector of *B* (blue) indicates collateral sensitivity. According to the position of the MIC and the breakpoints, the naive strain *V* 0 is sensitive to both drugs, the variant *V* 1 is resistant to both drugs and the variant *V* 2 is resistant to drug *B* but sensitive to drug *A*.

### 2.1 Defining sensitivity and resistance

Consider *N* drugs {*σ*_1_, *· · ·, σ*_*N*_ } = Σ. Σ defines a drug concentration space 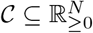, where *c* = (*c*_1_, *· · ·, c*_*N*_) *∈ 𝒞*is such that *c*_*i*_ represents concentration of drug *σ*_*i*_ for all *i* = 1, *· · ·, N* (see Figure 1 for 2 drugs Σ = {*A, B}*). We define a MIC *N* -dimensional vector that quantifies collateral sensitivity and cross-resistance of all drugs on Σ for a particular microorganism *V, MIC*(*V*) ∈ 𝒞, as follows

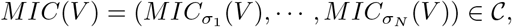

where 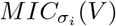 is the MIC of drug *σ*_*i*_ for microorganism *V*. The breakpoints of Σ, *Br*_Σ_ ∈𝒞, is given by

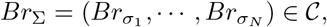

where 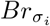 represents a maximum concentration of drug *σ*_*i*_ allowed to used for all *i* = 1, *· · ·, N*. Then, the breakpoints and the MIC of Σ state the following.

#### Definition 2

(Sensitive/Resistant strains). *Consider a microorganism V and N drugs* Σ = {*σ*_1_, *· · ·, σ*_*N*_ }. *It is said that V is sensitive/resistant to drug σ*_*i*_, *with i* = 1, *· · ·, N, if the i*^*th*^ *element of the vector MIC*(*V*) *− Br*_Σ_, *is negative/non-negative*.

The phenomenon of a microorganism *V*_*i*_ exposed over time to drug *σ* until it converges to a resistant variant *V*_*j*_ (a different phenotypic state) can be represented by the following graph

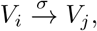

which can be measured by the following *drug-driven vector*.

#### Definition 3

(Drug-driven vector). *Consider the following graph* 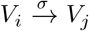. *The* drug-driven vector, 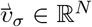, *is defined by*

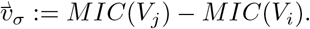

Negative elements of 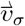 represent collateral sensitivity, unintendedly produced by drug *σ*; positive elements represent cross-resistance, unintendedly produced by drug *σ*, and null elements nonchange in relation to the MIC of the naive variant *V*_*i*_. In addition, considering the breakpoints of Σ and Definition 2, the vector 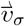 determines the susceptible/resistant profile of all emerging variants.

### 2.2 Evolutionary Network interactions between bacteria and antibiotics

To obtain a formal dynamical model describing the mutation/variation path of a pathogen when exposed to a sequence of *N* possible drugs (i.e., when applying one drug at a time), we associate each possible combination of sensible/resistant variant to a state in a state-space. This way, the potential dimension of the state space is 2^*N*^, accounting for the fact that each variant can be either sensitive or resistant to each of the drugs. Formally, an state *x*_*i*_, corresponding to variant *V*_*i*_, for some *i* ∈ {1, · · ·, 2^*N*^ }, is given (for instance) by *x*_*i*_ := (*σ*_1,*S*_, *σ*_2,*R*_, *· · ·, σ*_*N,S*_), meaning that variant *V*_*i*_ is susceptible to drug *σ*_1_, resistant to drug *σ*_2_, *· · ·*, and susceptible to drug *σ*_*N*_ or, the same, 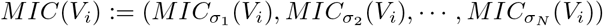 fulfill the conditions

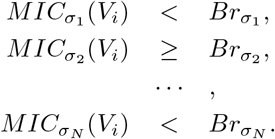

We denote the complete model containing the 2^*N*^ states as the classical one. However, if the model is intended to represent the system behavior just for realistic initial conditions - i.e., initial conditions that can be reached starting from other realistic initial conditions and feasible paths - the dimension (and complexity) of the model can be significantly reduced. Suppose that the original complete system of dimension 2^*N*^ is divided into two subspaces of dimension *L* and *E*, respectively (*L* denotes “latent” and *E* denotes ‘explicit’ subspace), such that *L* + *E* = 2^*N*^. And suppose also that the *L*-dimensional subspace cannot be reached, by any drug sequence, from its complementary subspace of dimension *E*, containing this latter all the realistic initial conditions. This way, the *E*-dimensional subspace is an invariant subspace containing all the initial biologically feasible conditions: *L*-dimensional subspace can not be the result of a mutation path promoted by the collateral sensitivity and cross-resistance profiles of the chosen drugs.

To exemplify this point, let us consider the hypothetical case of a pathogen and two drugs described in Figure 1, where drug *A* shows cross-resistance with respect to drug *B* and drug *B* presents collateral sensitivity with respect to drug *A*. According to this simple scheme, the complete 2^*N*^ states are denoted as *A*_*S*_*B*_*S*_, *A*_*R*_*B*_*S*_, *A*_*s*_*B*_*R*_ and *A*_*R*_*B*_*R*_, denoting all possible variant combinations according to their sensitivity/resistance to each drug. However, starting from the (wild type) variant *V*_0_, which is a *A*_*S*_*B*_*S*_ state, and a biologically feasible initial condition, only variants *V*_1_, which is *A*_*S*_*B*_*R*_, and variant *V*_2_, which is *A*_*R*_*B*_*R*_, can be reached by applying sequentially drugs *A* and *B* (in any order and during any time period). State variant *A*_*R*_*B*_*S*_ constitutes in this example the *L*-dimensional (with *L* = 1) latent subspace to which the system cannot evolve, while the complementary 3-dimensional subspace is the invariant sub-space. Figure 2 shows a graph 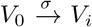 for every *σ* ∈ Σ := {*A, B*} and *i* = 1, 2, of this example. Our main hypothesis supporting this approach is that, given that the collateral sensitivity and cross-resistance profiles of the drugs are considered fixed for the model, they necessarily determine its structure (its dimension and potential behavior).

**Figure 2.**
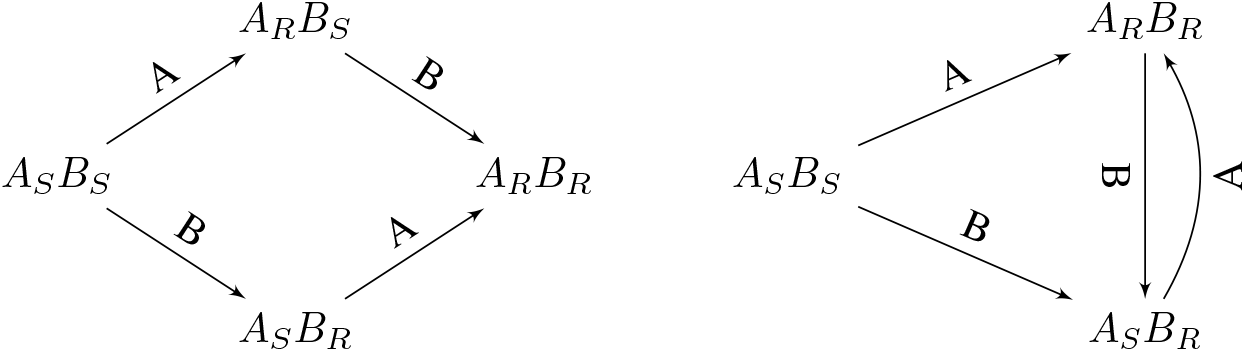
Classic combinatorial mutation network describing resistant and sensitive states for drugs *A* and *B* resulting in a total of 2^*N*^ states (left) vs a combinatorial mutation network accounting for collateral sensitivity and cross-resistance of drugs *A* and *B* (right).

The general way to obtain the reduced model is by applying each drug at a time to each state that emerges from the previous application of each drug. The procedure is stopped when the network converges to a closed graph, as shown in the right plot of Figure 2.

### 2.3 Dynamical model

A network with *n* sates (*x*_1_, *x*_2_, *· · ·, x*_*n*_) can be linked to a switched system, as demonstrated in a prior study [61] and by this we can investigate the impact of sequential drug exposures on population dynamics. The effect of therapy is modeled by drug-induced death rate framework: the growth rate for variant *x*_*i*_ under exposure of drug *σ* is given by 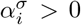 and death rate 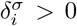. Therapy (*σ*) can affect also mutation from *x*_*i*_ to *x*_*j*_ rates by 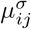. The combinatorial mutation network is given by the binary matrix 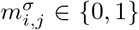, where 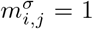 implies an arrow from *x*_*i*_ to *x*_*j*_, and 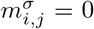 there is no connection. The total carrying capacity is *K >* 0. The following switched system describes the nonlinear dynamics between states.

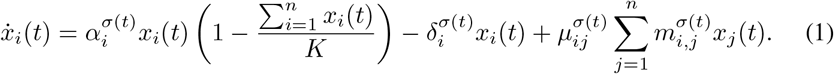

The states of the system are given by *x* = (*x*_1_, *x*_2_, *· · ·, x*_*n*_), and the control input by *σ*, for all time *t >* 0.

#### Remark 1.

*For switched systems, mode selection from the finite set can be formulated as a dynamic programming problem, but the solution is difficult to implement and may not account for constraints [67, 59]. More applicable control approaches involve mixed-integer optimization [68], state-dependent switching rules [69], and receding horizon switching signals [58]*.

The switch of drugs shifts the balance of birth and death such that the colony shrinks or escapes. It is assumed that during single-drug therapy with, say, drug *σ*_*d*_, mutations lead to the emergence of a variant resistant to this drug, i.e., state *σ*_1,?_, *· · ·, σ*_*d,S*_, *· · ·, σ*_*N*,?_ varies/mutates to state *σ*_1,?_, *· · ·, σ*_*d,R*_, *· · ·, σ*_*N*,?_ whatever ? is *S* or *R*. Consequently, the resistant phenotype proliferates while the sensitive strains will be killed at a constant rate, ultimately culminating in treatment failure. Also, to simplify the model, all sensitive cells are killed at the same rate as the naive strain.

#### Remark 2.

*The dynamics framework we consider here is that under the pressure of any drug there is an evolution towards resistance to the drug. Thus monotherapy eventually fails to contain overall infection levels leading to treatment failure as resistant strains dominate. Within this framework, combination drug regimens are required to prevent uncontrolled spread of the infection. Our goal is to determine optimal switching protocols that constrain the infection despite ongoing evolution of drug resistance*.

## 3 Invariance analysis for drug-resistant infections

Set-control theory deals with the properties of some regions of the state space that are applicable to both the practical and theoretical aspects of control [54]. Particularly, the invariant sets of a dynamical system are regions of the state space that trap the trajectories of every state entering them: once the system is inside, it cannot escape the invariant set. In the following, we detail how the theory of invariant sets can be applied to analyze bacterial resistance.

In the framework proposed by Eq. (1) it is required to use combinations of antibiotics with mutual collateral sensitivity to prevent population trajectory to escape, because of the constant evolution towards resistance. However, this requirement is not a sufficient condition to ensure that population trajectories will not escape. To see this, consider Figure 3, where three illustrative dynamics show the effects of drug *A* (red) and drug *B* (blue) with mutual collateral sensitivities on the two populations *A*_*R*_*B*_*S*_ and *A*_*R*_*B*_*S*_. Depending on the birth, death, and mutation rates, (a) the behavior of the population can achieve perfect balance and the total population can be maintained inside a target window; (b) the population may experience shrinkage so it will remain inside the target window; and (c) the population may escape, because it is not possible to keep the states inside the target region.

**Figure 3.**
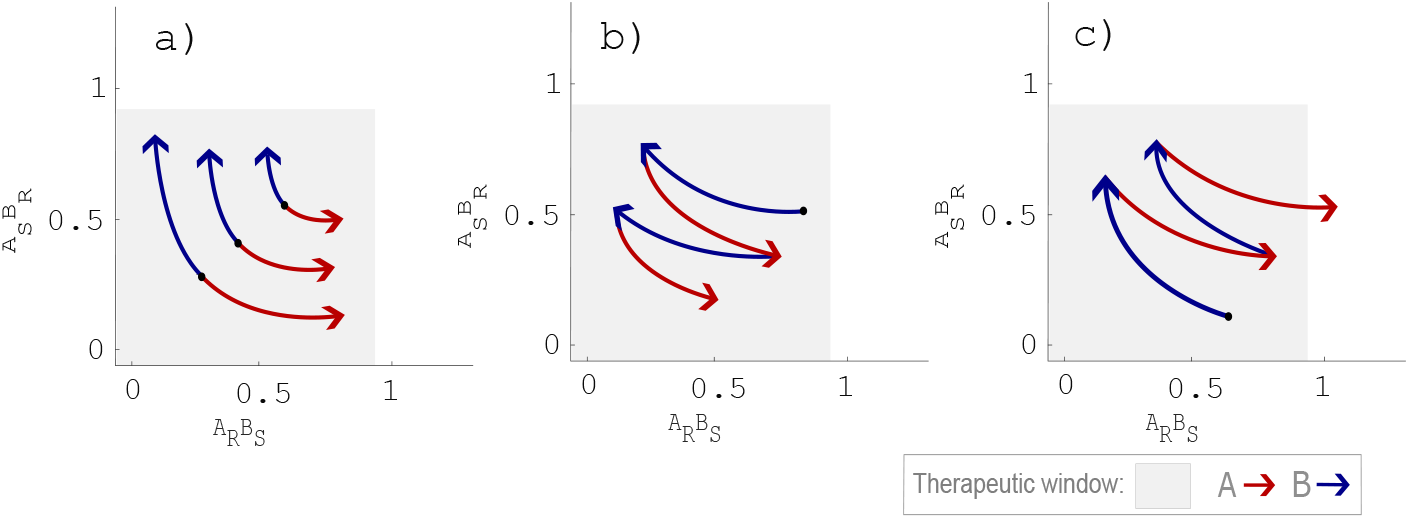
**Illustrative dynamic evolution of mutual collateral sensitivity**. Exposure to drug *A* reduces concentration of *A*_*S*_*B*_*R*_ cells (sensitive to *A* and resistant to *B*) but allows the escape of *A*_*R*_*B*_*S*_. Oppositely, exposure to drug *B* reduces concentration of *A*_*R*_*B*_*S*_ but induces the escape of *A*_*S*_*B*_*R*_. a) A proper balance between dynamic of drugs *A* and *B* implies that the total population can be feasible bounded. b) The best scenario of mutual collateral sensitivity implies the asymptotic stability of the origin. c) Despite a mutual collateral sensitivity it may not be feasible to prevent the escape of the infection.

The assessment of this problem has been successfully tackled by set-control theory [54]. Set-control involves examining regions within the state space that feasibly bound system trajectories; such regions are known as invariant sets. Examining invariant sets for our dynamical system can help us to determine if a trajectory will remain bounded or escape from a target window..

Let us assume there is a therapeutic window 𝕋 on the states space (containing the origin) where we want to maintain the population of microorganisms. We consider a discretization of the system (1) for a given step size. A control invariant set (CIS) inside 𝕋, for the discrete switched system can be defined as follows.

### Definition 4

(Control Invariant Set - CIS). *A set* Ω *⊂* 𝕋 *on the states space is said to be a control invariant set of the switched system if for every state x*(*k*) ∈ Ω *there is a feasible mode σ* ∈ Σ *such that the next state follows the condition x*(*k* + 1) ∈ Ω.

The invariant set is a generalization of the steady-state (equilibrium) concept and its existence implies there is a feasible sequence of drugs that keep the population bounded inside Ω, obviating so far the need for identifying the specific drug sequence, its order and exposure times, which is related with a complex and challenging optimal control problem. Instead, by simply ensuring the existence of a CIS, we can establish that the infection can be feasibly contained. Certainly, the presence of a multidrug-resistant (MDR) strain within the dynamical network implies the nonexistence of an invariant set within a therapeutic window, provided the latter is bounded to safety regions.

To characterize a CIS, the concept of controllable set can be used [58]. The controllable set of Ω is given by all the states that can be driven by a feasible mode *σ* ∈ Σ in one step to Ω, and it can be formally defined as follows:

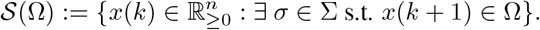

The next result is well established on set-control theory.

### Property 1.

*If* Ω *⊆ 𝒮* (Ω) *then* Ω *is a CIS*.

*Proof*. Suppose that *x*(*k*) ∈ Ω for some time-step *k ≥* 0. Since Ω *⊆ 𝒮* (Ω), then *x*(*k*) ∈ 𝒮 (Ω) as well. By definition of the controllable set 𝒮(Ω) there exists *σ* ∈ Σ such that *x*(*k* + 1) *∈* Ω, which conclude the proof.

Various methods for characterizing invariant sets for a linear switched system were proposed in [55]. Based on the ideas presented there, we have developed a method for characterizing CIS in our framework, as detailed in the next section.

### 3.1 Invariant sets characterization

For several infectious diseases, nutrients are in excess. Thus, the nonlinear model described by Equation (1) can be simplified to a linear model when 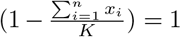. In addition, if we consider mutation rate 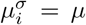 for all *i* = 1, *· · ·, n* and *σ ∈* Σ, Eq. (1) can be rewritten as follows:

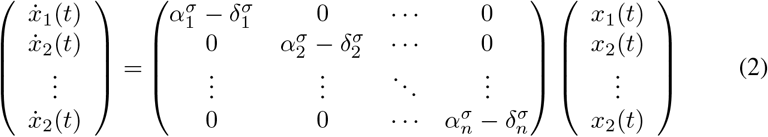

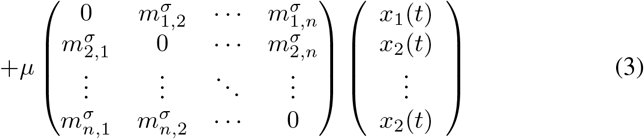

which is equivalent to the switched linear system given by the following equations

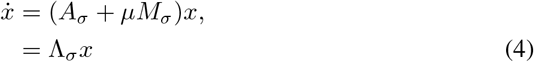

where *x* belong to the state constraint 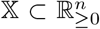 and *σ ∈* Σ. Note that 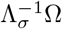 is well defined even if matrix ?_*σ*_ is singular.

#### Lemma 2.

*A set* Ω *⊂* 𝕏 *is a control invariant set for the switched linear system* (4) *if and only if* 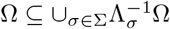.

We propose the following algorithm to compute a control invariant for the linear switching system within a window 𝕋 *⊂* 𝕏.

#### Algorithm 1

Invariant set within 𝕋 for *N* drugs

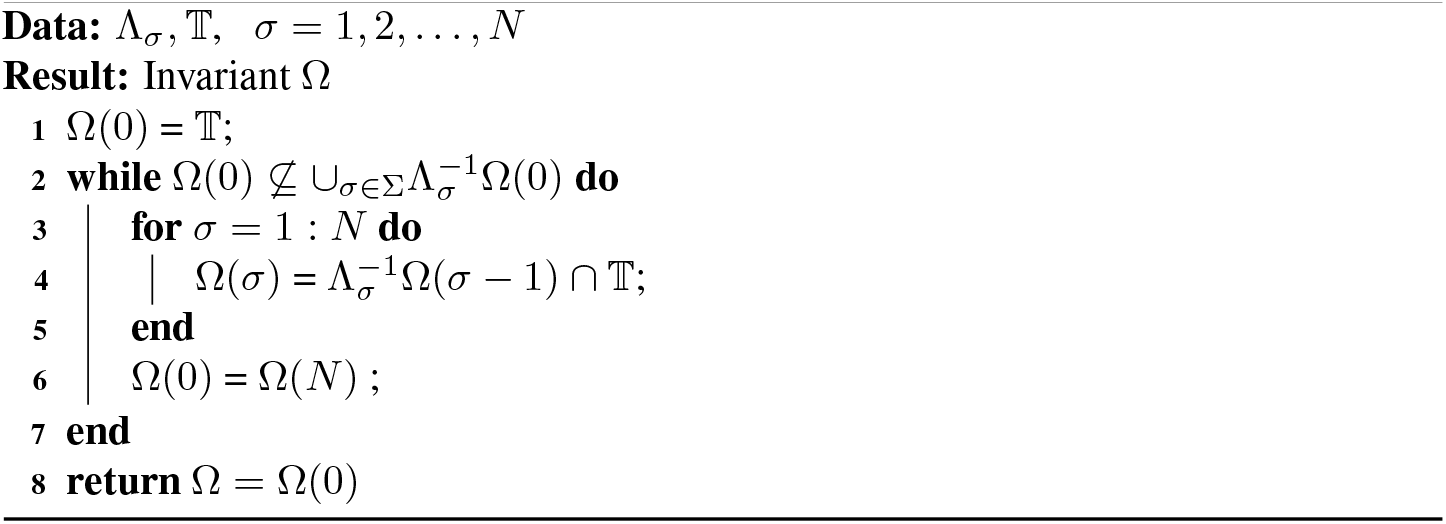

#### Proposition 3.

*Algorithm 1 converges to the largest invariant set of the switched system* (4) *contained in* 𝕋, *or to an empty set*.

## 4 Numerical Results

We assessed methods to find bounded invariant sets for the switched system associated with different treatments, and we demonstrate that the existence of this set implies the infection can be suppressed (or at least contained) through a feasible sequence of drugs. We use data collected in Imamovic et al. for *P. aeruginosa* strain PAO1 exposed to clinically significant antibiotics, increasing the concentration over 10 days such that by day 10 all strains could grow in the presence of the antibiotic when the antibiotic concentration was above the EUCAST breakpoint. Dose response curves were performed with the 23 other antibiotics to elucidate collateral sensitivity or collateral resistance interactions showed in the heatmap on Figure 4.

**Figure 4.**
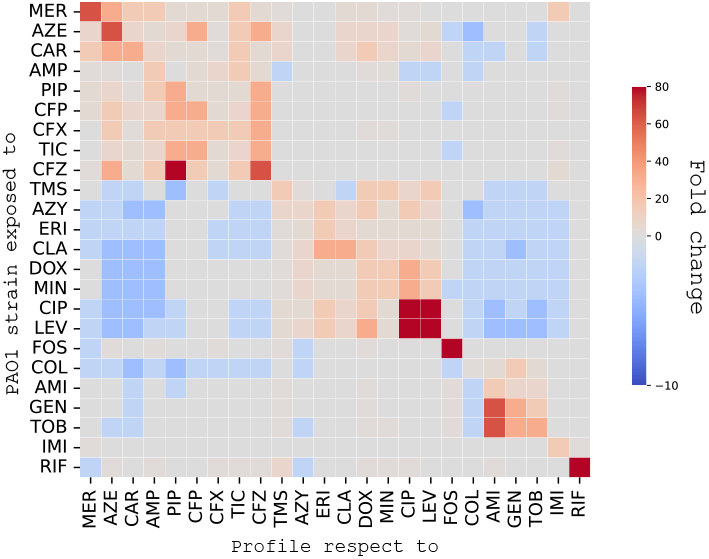
Heat-map representation of collateral sensitivity and resistance from [29]. Antibiotics are specified on Table A1.

### 4.1 Sequential drug therapy simulation

We have simulated a combinatorial mutation network to describe the interactions between strains resistant and sensitive to 3 drugs. We compare a cyclic treatment for two types of networks. The first is a conventional network with 3 generic drugs *A, B* and *C*; in this case there are 2^3^ different phenotypic states (as described in Figure 5-a), the drug-driven resistance leads to mutations that follow a dead-end path to the multidrug resistance strain *A*_*R*_*B*_*R*_*C*_*R*_, which is inevitable. The second network is designed according methodology proposed on Section 2.2, for the antibiotics Aztreonam (AZE), Ciprofloxacin (CIP), Tobramycin (TOB) from Table A1, with *A* = *AZE, B* = *CIP*, and *C* = *TOB*. The collateral sensitivity and cross-resistance profiles of these drugs according Figure 4 (generated in [29]) produce a particular network, where the directions of mutations take into account all the side effects. Figure 5-b shows the network for these drugs, the red arrows indicate that there is cross-resistance and the blue arrows indicate that there is collateral sensitivity. As there are 3 drugs, one drug can exhibit cross-resistance concerning one drug and collateral sensitivity concerning the other; in these cases the black arrow is used. Both networks were linked to the dynamics model in Equation 1. This allowed simulating bacterial growth under 50*hr* cyclic dosing of drugs *A, B, C* over 600*hr* (as described in Figure 5-c). Importantly, Figure 5-b shows that drug interactions alter potential mutation pathways, so not all theoretically possible phenotype strains manifest in the network. The bacteria cannot mutate in a way that becomes fully resistant to all drugs, what cannot be avoided in the classic network presented in Figure 5-a. This is a crucial point as it enables us to select the right drugs to prevent the emergence of multiresistant strains.

**Figure 5.**
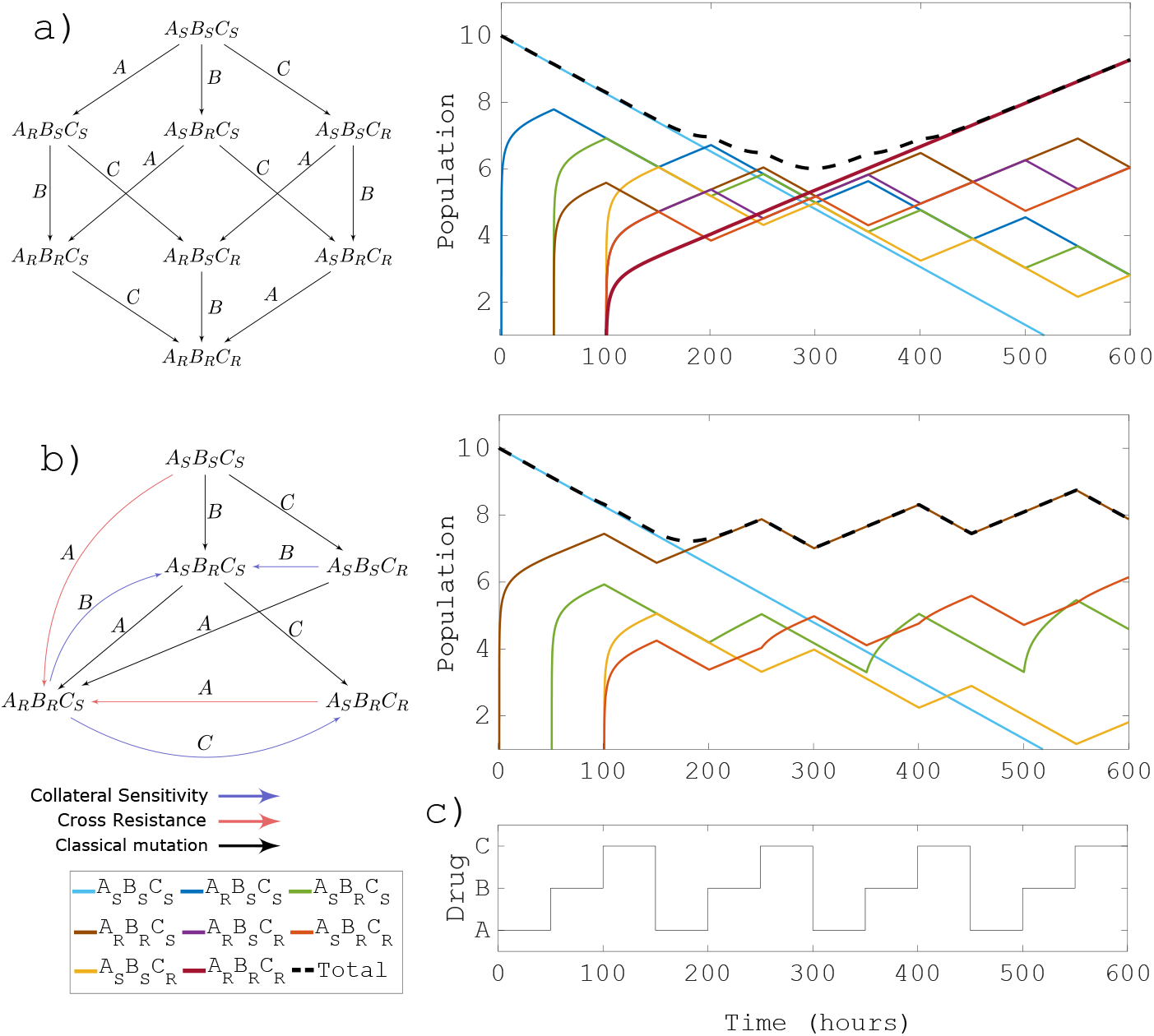
Dynamic evolution of a classic mutation network vs a collateral sensitivity network simulations for three drugs with cyclic treatment. We consider two cases: a) for a combinatorial mutation network given by generic drugs without collateral effects (a). And b) for a combinatorial mutation network that considers collateral effects between drugs Aztreonam (*A* = *AZE*), Ciprofloxacin (*B* = *CIP*), Tobramycin (*C* = *TOB*).

### 4.2 Invariant set for Levofloxacin and Ampicillin

The antibiotics Levofloxacin and Ampicillin show mutual collateral sensitivity when used in PAO1 wild type [29], thus providing an ideal scenario to test whether the model generated by these two antibiotics admit invariants within a therapeutic window. The mutual collateral sensitivity of the drugs LEV and AMP generates the combinatorial mutation network shown in Figure 6-a, with the states *A*_*S*_*B*_*S*_ (sensitive to both drugs), and the states *A*_*S*_*B*_*R*_ and *A*_*R*_*B*_*S*_. The network indicates the direction of mutations under the effect of each drug. Note that the system has 3 states and one of them is sensitive to both drugs (wild type *A*_*S*_*B*_*S*_), therefore this state converges to the origin when either drug *A* or drug *B* is applied. Thus, we can disregard this dimension, and represent the invariant sets in 2 dimensions; *A*_*S*_*B*_*R*_ and *A*_*R*_*B*_*S*_. In this way, we have better visibility of the sets. To calculate a control invariant for the system associated with the network, we use Algorithm 1. Four cases were simulated, in which different growth, death, and mutation rates were considered (see Section 5). For cases one (Figure 6-b), two (Figure 6-c), and four (Figure 6-e) we chose a therapeutic window given by the set 𝕋_1_ = {0 *≤ A*_*R*_*B*_*S*_ *≤* 1, 0 *≤ A*_*S*_*B*_*R*_ *≤* 1}, and the algorithm finds an invariant within 𝕋_1_, as Figure 6 shows the controllable sets contains set Ω in each case, fulfilling Property 1. For case 3 (Figure 6-d) the algorithm does not find an invariant within 𝕋_1_, but it converges for 𝕋_2_ = {0 *≤ A*_*R*_*B*_*S*_ *≤* 1, 0 *≤ A*_*S*_*B*_*R*_ *≤* 0.5}.

**Figure 6.**
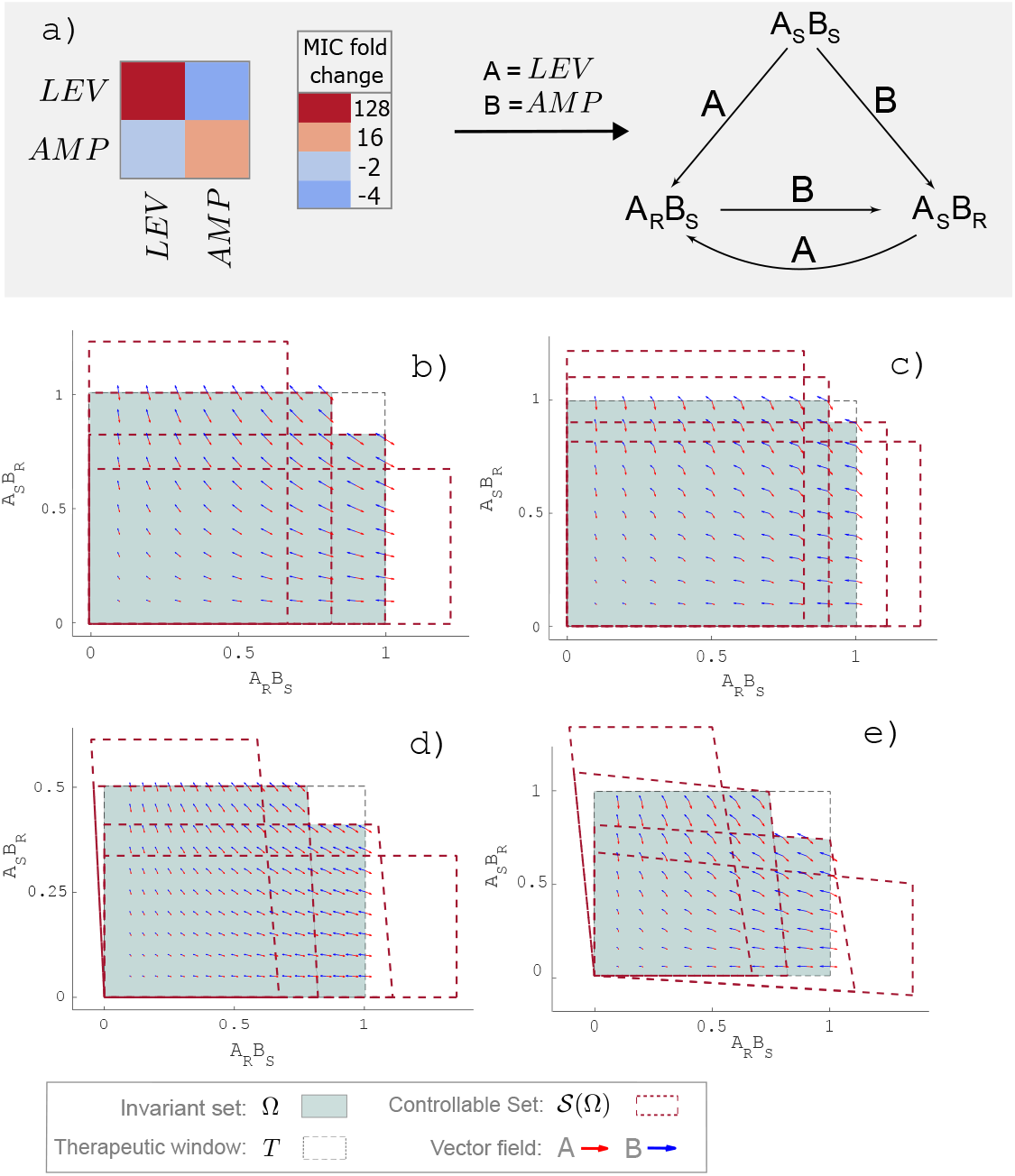
Invariant set for combination of drugs Levofloxacin (*LEV*) and Ampicillin (*AMP*). From the profiles of collateral sensitivity and cross resistance of drugs *LEV* and *AMP* the network describing the interactions between variants *LEV*_*S*_*AMP*_*S*_, *LEV*_*R*_*AMPC*_*S*_ and *LEV*_*S*_*AMPC*_*R*_ is computed. There exists invariant sets of the mathematical model associated to the network which implies the infection can be bounded with drugs *LEV* and *AMP*. Invariant Set Ω (green) inside *T*. Note that the controllable set 𝒮 (Ω) (dashed-lines) contains the invariant set for all cases.

### 4.3 Invariant set for Tobramycin, Carbenicillin and Colistin

Here we consider 3 drugs; Tobramycin, Carbenicillin, and Colistin, whose collateral effects were analyzed in [29] for the treatment of the PAO1 wild-type strain. According to Imamovic et. at. these drugs exhibit collateral sensitivity and cross-resistance both. The combinatorial mutation network for these drugs is shown in Figure 7-a, with 4 states: the wild type *A*_*S*_*B*_*S*_*C*_*S*_ sensitive to all drugs (*A* = *T OB, B* = *CAR, C* = *COL*), and three other states that mutate among themselves according the collateral effects of the drug used, *A*_*S*_*B*_*R*_*C*_*S*_, *A*_*R*_*B*_*S*_*C*_*S*_ and *A*_*R*_*B*_*S*_*C*_*R*_.

**Figure 7.**
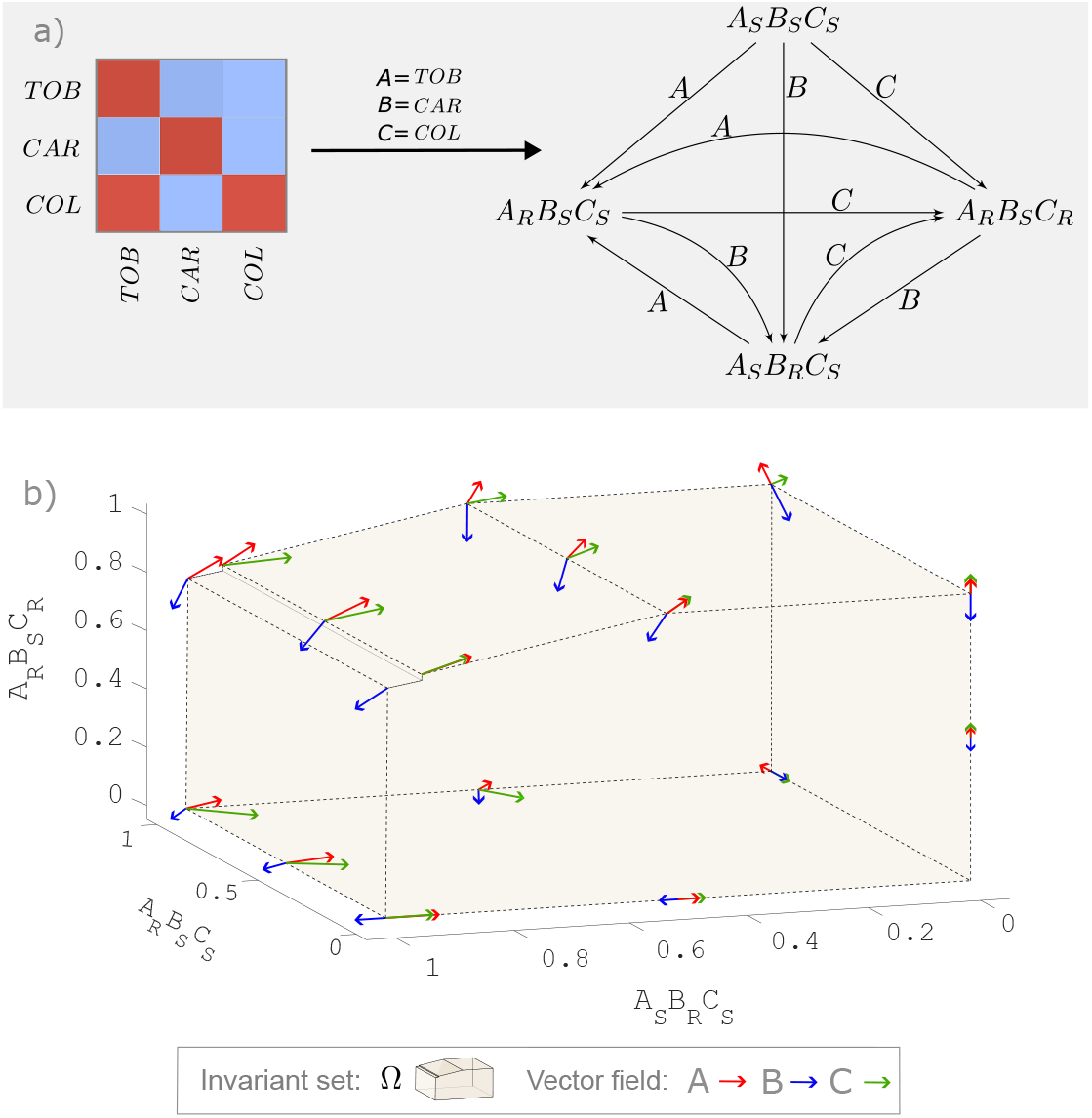
Normalized cell population of variants *A*_*R*_*B*_*S*_*C*_*R*_, *A*_*R*_*B*_*S*_*C*_*S*_ and *A*_*S*_*B*_*R*_*C*_*S*_ inside the invariant set Ω for drugs Tobramycin (*TOB*), Carbenicillin (*CAR*) and Colistin (*COL*).

Through Algorithm 1 we proved that there exists an invariant for the linear dynamic system associated with this set of drugs, which is plotted in Figure 7-b in three dimensions (the dimension of *A*_*S*_*B*_*S*_*C*_*S*_ is not considered because this state always decays), the parameters from Table 3 were used.

The invariant set Ω found and represented in Figure 7-b implies that the infection can be feasibly contained through some sequence given by the drugs *T OB, CAR* and *COL*. This means that even though there is constant evolution towards resistance, the dynamics and interaction between these drugs are enough to contain the infection. Technically, if a state,

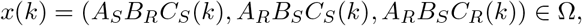

representing population at time *k ≥* 0, one drug in Σ = {*T OB, CAR, COL}* implies that

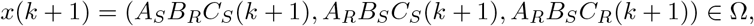

and so on indefinitely.

## 5 Discussion

Our finding shows that a sequential drug treatment that aims to eliminate a population of distinct bacterial phenotype all evolving resistance to the current drug requires drug synergy. Without collateral sensitivity between agents, single or combined antibiotics are bound to fail by enabling persistence of at least one resistant bacterial variant. In this current study, we have used a birth/death model that captures interactions between distinct variants of bacteria using the collateral sensitivity and cross-resistance profile of the drugs targeting specific species. The interactions were modeled by combinatorial mutation networks where the mutation rates between the variants can be specified.

We modeled the drug effect by discrete events affecting the birth, death, and mutation rates of a continuous bacterial population. Systems containing both continuous and discrete-time dynamics and characterized by abrupt changes in at least one state variable at certain time instants, are called hybrid systems [70] and can be used to analyze qualitative properties of the population evolution.

Our model enables predicting the effect of sequential drug treatments, taking into account the antagonistic and synergistic interactions between the drugs involved in the process. This proposed model can be used to study advanced control strategies and determine optimal combinations and sequences of drugs. It can also be used to determine which combinations lead to the proliferation of multidrug-resistant strains, as well as to analyze the trade-offs between the ongoing evolution of resistance to the current drug and the evolution toward sensitivity to potential drugs. In the latter, we focused some of our dynamic analysis. To this end, we employed set-control theory to characterize stable regions within the feasible state space. These regions could include steady states, multiple equilibria, or invariant sets. However, our system had only healthy equilibria (e.g., the origin) with no relevance in this context. Therefore, we proposed to use the concept of control invariant sets to assess when the total population could be feasibly maintained within a therapeutic window, as an invariant set captured the regions of the state space where the system can feasibly remain indeterminately. We provided a method to characterize invariant sets for our particular system and computed several invariant sets for a population of distinct phenotypes of P. aeruginosa and its sensitivity profile for 24 relevant antibiotics [29]. In this way, the existence of a nonempty invariant set within a therapeutic window for a specific system (determined by the drugs chosen for therapy) ensures that drug resistance could be feasibly contained. Furthermore, the nonexistance of such a set implies that the population can not be feasibly contained inside the therapeutic window, resulting in resistance prevailing over synergy.

Our dynamic analyses offer valuable qualitative and quantitative insights into the collateral effects on population dynamics. We can reliably identify drug combinations leading to treatment failure and those enabling the controlled dynamics of all subpopulations, depending on specific birth, death, and mutation rates. To expand our findings, we must incorporate PD/PK factors, account for stochastic processes, address resource competition, and delve into parameter estimation. Although these additional factors introduce a layer of complexity to the dynamic analysis, also would allow us to effectively leverage our results in modeling in-vitro population growth, and to ensure the effectiveness of combining pharmacology and evolution for sustaining pathogen suppression.

### Parameters and software availability

For parameter simulations, we maintained uniform values for mutation rate and carrying capacity across all cases. Specifically, we set 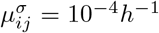 for all *i* = 1, *· · ·, n* and for all *σ ∈* Σ. Additionally, we adopt qualitative values for growth and clearance rates, such that 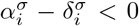 for strains *x*_*i*_ sensitive to drug *σ* and 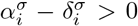 for strains *x*_*i*_ resistant to drug *σ*. Finally, only for the analysis of Figure 6, extreme values (on the order of the growth rate) of the mutation rate were considered in order to obtain qualitative results.

Software is available at https://github.com/AlejandroRedna/Invariant_Bacterial_Resistance_Mathematical_Bioscience_2024.

## Acknowledgment of Support and Disclaimer

This material is based upon work supported by the National Science Foundation under Grant No. 2315862.

## Appendix

**Table A1:** List of antibiotics used in [29] for the evolution of wild-type P. aeruginosa.

| ANTIBIOTIC | ABBREVIATION | CLASS | TARGET |
| --- | --- | --- | --- |
| Amikacin | AMI | aminoglycoside | protein synthesis, 30S |
| Gentamicin | GEN | aminoglycoside | protein synthesis, 30S |
| Tobramycin | TOB | aminoglycoside | protein synthesis, 30S |
| Ciprofloxacin | CIP | quinolone | DNA gyrase |
| Levofloxacin | LEV | quinolone | DNA gyrase |
| Ampicillin | AMP | b-lactam (penicillin) | cell wall |
| Piperacillin | PIP | b-lactam (penicillin) | cell wall |
| Carbenicillin | CAR | b-lactam (penicillin) | cell wall |
| Ticarcillin | TIC | b-lactam (penicillin) | cell wall |
| Aztreonam | AZE | b-lactam (monobactam) | cell wall |
| Cefepime | CFP | b-lactam (cephalosporin) | cell wall |
| Cefuroxime | CFX | b-lactam (cephalosporin) | cell wall |
| Ceftazidime | CFZ | b-lactam (cephalosporin) | cell wall |
| Meropenem | MER | b-lactam (carbapenem) | cell wall |
| Imipenem | IMI | b-lactam (carbapenem) | cell wall |
| Minocycline | MIN | tetracycline | protein synthesis, 30S |
| Doxycycline | DOX | tetracycline | protein synthesis, 30S |
| Azithromycin | AZY | macrolide | protein synthesis, 50S |
| Erythromycin | ERI | macrolide | protein synthesis, 50S |
| Clarithromycin | CLA | macrolide | protein synthesis, 50S |
| Colistin | COL | polymyxin | lipopolysaccharide |
| Fosfomycin | FOS | fosfomycin | cell wall biogenesis |
| Rifampicin | RIF | rifamycin | RNA synthesis |
| Trimethoprim | TMS | antifolate | combination folic acid pathway |

